# Influenza-induced activation of recruited alveolar macrophages during the early inflammatory phase drives lung injury and lethality

**DOI:** 10.1101/2020.06.08.141309

**Authors:** Clarissa M Koch, Kishore R Anekalla, Yuan-Shih Hu, Jennifer M. Davis, Mark Ciesielski, Gaurav Gadhvi, Shang-Yang Chen, Margaret Turner, Yuan Cheng, Bria M Coates, Hiam Abdala-Valencia, Paul A Reyfman, Alexander V Misharin, GR Scott Budinger, Deborah R Winter, Karen M Ridge

## Abstract

Severe respiratory virus infections initiate a robust host immune response that contributes to disease severity. Immunomodulatory strategies that limit virus-initiated inflammation are of critical importance. In this study, we compared the host response to influenza A virus (IAV) infection in susceptible animals (wild-type, WT) to resilient mice (*Vimentin^-/-^* mice). We identified distinct gene expression patterns in recruited monocyte-derived alveolar macrophages (MoAMs) associated with three phases (Infiltrating, Early Inflammatory, Late Inflammatory) that evolve in sequence over the course of IAV infection. We report a core set of pro-inflammatory genes involved in the WT MoAM Early Inflammatory response that is suppressed in *Vim^-/-^* MoAMs. Moreover, we identify CEBPB, Jun-AP1, and IRF transcriptions factors as regulators of this attenuated inflammatory response. We performed causal experiments using bone-marrow chimeras to credential that *Vim^-/-^* MoAMs with suppressed pro-inflammatory genes confer protection from influenza-induced mortality in WT susceptible mice. Taken together, these data support the notion that vimentin plays a causal role in determining the pro-inflammatory function of recruited MoAMs and drives IAV-induced lung injury.

## Introduction

Severe viral pneumonia due to influenza A virus (IAV) or severe acute respiratory syndrome coronavirus 2 (SARS-CoV-2) is characterized by the release of proinflammatory mediators that damage the alveolar epithelial and capillary barrier to cause acute respiratory distress syndrome (ARDS)[1, 2]. This points to the importance of a balanced and regulated immune response, where infection can be controlled while keeping tissue damage to a minimum. This hypothesis is clinically supported by a recent analysis of patients enrolled in the ARDSnet, in which patients with a hyperinflammatory endotype of ARDS, characterized in part by increased levels of circulating cytokines and chemokines, had worse clinical outcomes [3].

Shortly after IAV or SARS-CoV-2 infection, tissue-resident alveolar macrophages (TRAMs) are depleted, and monocytes are released from the bone marrow and recruited into the infected, injured lung [1, 4–6]. Upon entering the lung, these recruited monocytes, referred to as monocyte-derived alveolar macrophages (MoAMs), drive the immune response in humans and mice. MoAMs activate inflammatory signaling pathways in response to pathogen- or damage-associated molecular patterns (PAMPs or DAMPs) released from infected and dying cells [1, 3, 4, 7]. Preventing the recruitment of MoAMs into the lung by deleting CCR2 or its ligand reduces the severity of the dysregulated immune response (e.g., no cytokine storm) without affecting viral clearance [4]. These data suggest that recruited MoAMs mediate the excessive pro-inflammatory response associated with tissue damage. Indeed, elevated levels of IL-1β and IL-18, cytokines produced by the NLRP3 inflammasome, in the alveolar exudate of IAV-infected patients serve as poor prognostic markers of viral pneumonia outcomes [8–10]. Inhibiting the NLRP3 inflammasome early in IAV infection impairs viral clearance and increases morbidity and mortality in mice [11]. In contrast, inhibiting the NLRP3 inflammasome post-viral clearance reduces inflammation-induced tissue damage and promotes lung repair [11, 12]. These studies highlight the importance of understanding that recruited alveolar macrophages may play different roles in the virus-infected lung depending on the stage of infection (e.g., pre-versus post-viral clearance). As a result, therapies that target recruited MoAMs may be of interest [4, 13, 14]; however, no study has yet investigated the time-dependent transcriptional signature and function of MoAMs in response to IAV infection.

Vimentin, a type III intermediate filament, is ubiquitously expressed in macrophages and plays a role in several critical immune response processes. Vimentin expression is increased in COVID-19 patients’ lungs compared with healthy, control and COPD patients [15]. In patients with ulcerative colitis and Crohn’s disease, vimentin expression was also significantly increased in inflamed tissues [16]. Consistent with these observations, vimentin-null (*Vim^-/-^*) mice develop significantly less gut inflammation compared to wild-type mice with acute colitis induced by dextran sodium sulfate [17]. We have shown that *Vim^-/-^* mice are protected from LPS-induced lung injury and death by suppressing pro-inflammatory cytokines [18]. Vimentin is a known ligand for some PRRs, including nucleotide-binding oligomerization domain-containing protein 2 (NOD2) and NLR Family Pyrin Domain Containing 3 (NLRP3). We, and others, reported that vimentin is required to activate the NLRP3 inflammasome [18, 19] and subsequent NF-κB activation [20], which is consistent with the observation that pro-inflammatory gene expression is altered in microglia lacking vimentin in a murine model of Alzheimer’s disease [21]. Interestingly, bulk RNA-seq transcriptomic analysis showed significant upregulation of *Vim*, which was associated with decreased viral host resistance in cultured embryonic stem cells and embryos [22]. Collectively, these studies support the notion that vimentin may play a role in the immune response to IAV infection.

The temporal transcriptional response of recruited MoAMs that drives the immune response in the IAV-infected host has not been previously reported. For the first time, our data identifies the distinct gene expression patterns in recruited MoAMs associated with three phases (Infiltrating, Early Inflammatory, Late Inflammatory) that evolve in sequence following infection with influenza A virus. Additionally, we report that wild-type mice mount a sustained pro-inflammatory response contributing to increased lung injury and death. In contrast, *Vim^-/-^* mice are remarkably resilient to IAV infection. Whether vimentin plays a causal role in determining the pro-inflammatory function of recruited MoAMs in the IAV-infected lung has not been previously investigated. If vimentin indeed affects MoAMs’ pro-inflammatory function, is it a cell-autonomous phenomenon? Using isochronic adoptive MoAM transfer experiments and molecular profiling in mice, we sought to determine whether the loss of vimentin-related protection from influenza-induced lung injury is intrinsic to *Vim^-/-^*MoAMs. Our data supports a paradigm in which recruited MoAMs upregulate vimentin expression, activate maladaptive responses, and consequently exhibit a persistent pro-inflammatory response, promoting virus-induced lung injury.

## Results

### Vimentin null mice are protected from influenza-induced lung injury and mortality

Our group has recently reported that persistent inflammation in patients with severe SARS-CoV-2 pneumonia is sustained via stable circuits between recruited alveolar macrophages and T cells[1]. Using single-cell RNA-seq, distinct clusters of monocytes-derived alveolar macrophages (MoAMs) were identified from the bronchoalveolar lavage fluid (BALF) collected from patients with severe SARS-CoV-2 pneumonia (**Figure 1a, Supplemental Figure 1a**) [1]. MoAM1 cluster was characterized by high expression of pro-inflammatory cytokine genes, including *IL1B*, *CXCL2*, and *CCL3*, and low expression of reparative genes *MRC1* and *C1QA* (**Figure 1c, d**) [23–25]. In contrast, cluster MoAM3 exhibited the highest expression of *MRC1*, *C1QA*, *and MERTK*. *MERTK* encodes a receptor necessary for phagocytosis of apoptotic cells – the uptake of apoptotic cells reduces the expression of pro-inflammatory cytokines and chemokines from macrophages. This process has been mechanistically linked to transcriptional responses leading to a change in macrophage phenotype from pro-inflammatory to reparative [26, 27] (**Figure 1c-d**). By querying this single-cell RNA-seq dataset, we found that *VIM* was among the genes differentially expressed between MoAM1 and MoAM3: with the highest expression in the pro-inflammatory MoAM1 population and decreasing in the reparative MoAM3 population (**Figure 1b-c**). These data from patients with severe SARS-CoV-2 pneumonia show that higher *VIM* expression is associated with a pro-inflammatory MoAM phenotype in viral pneumonia, while lower levels of *VIM* expression characterize MoAMs with a reparative phenotype. Importantly, these data suggest that there may be a causal relationship between *VIM* expression and MoAM phenotype.

**Figure 1.**
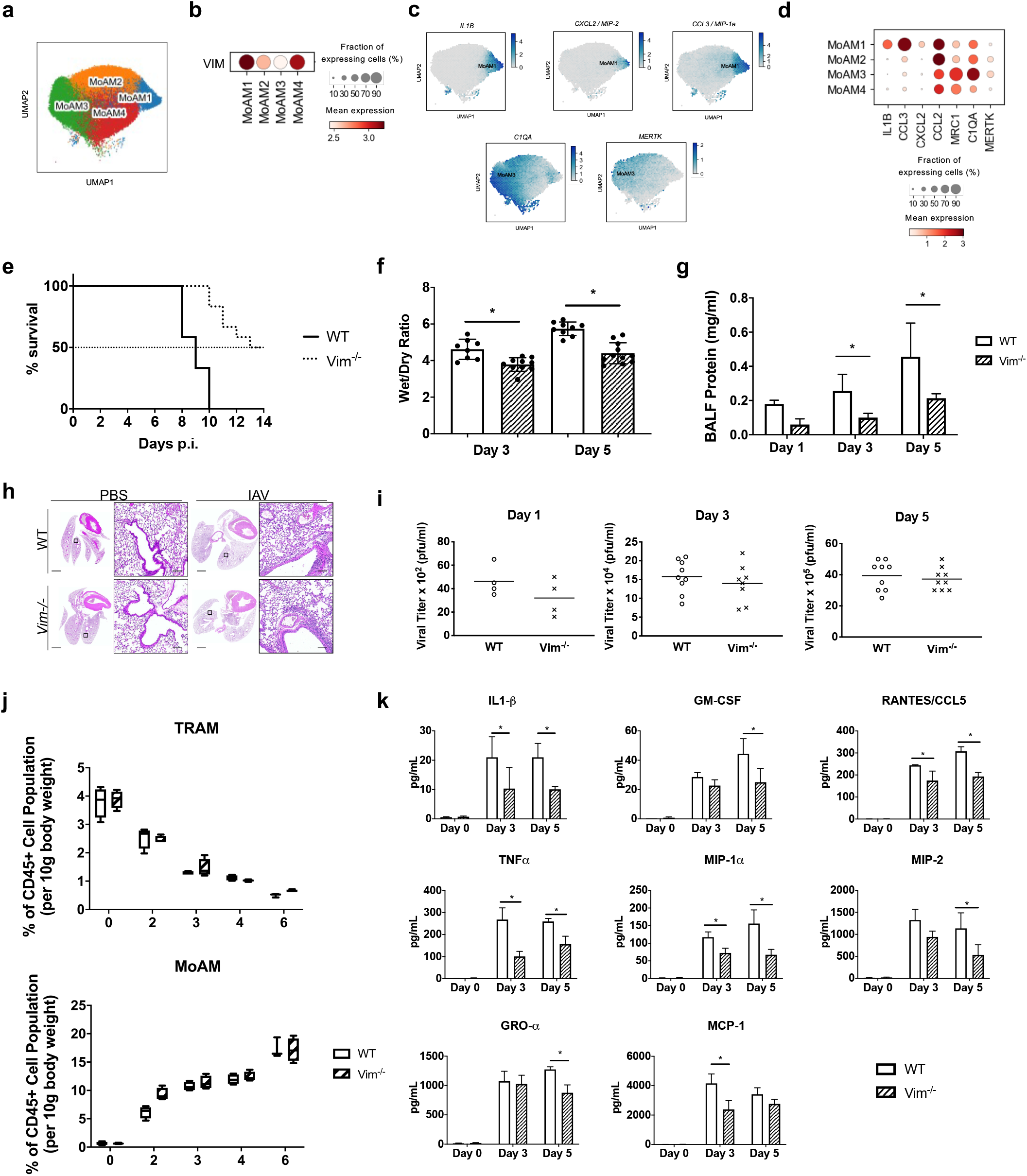
Vimentin-null mice are protected from influenza-induced lung injury and mortality. **a**) Uniform Manifold Approximation and Projection (UMAP) of 40,248 cells annotated as monocyte-derived alveolar macrophages (MoAMs) obtained from 10 patients with severe COVID-19 disease within 48 hours after intubation. **b**) Fraction of cells expressing *VIM* and mean expression in each MoAM cluster. **c**) Visualization of *IL1-B*, *CCL3*, *CXCL2*, *C1QA*, and *MERTK* expression in MoAM cells using density projection plots. Scale bar represents expression averaged within hexagonal areas on UMAP. **d**) Fraction of cells expressing the representative genes and their mean expression in each MoAM cluster. **e**) Survival of wild-type (WT) and *Vim^-/-^* mice intratracheally infected with a lethal dose of influenza A virus (IAV) (200pfu) was measured and compared. Survival curves were analyzed using log-rank test, p-value =0.001. Data shown are n=12 per group. **f**) Lung wet-to-dry ratio measured at day 3 and 5 p.i. (n = 8-11/timepoint) **g**) Bronchoalveolar lavage fluid (BALF) protein levels measured by Bradford assay, n=5/timepoint. In **f** and **g**, differences between genotypes on each day were assessed by t-test. p-values were 0.0015 and <0.0001, for 3 and 5 days p.i., respectively. **h**) Representative images of murine lung architecture at day 7 p.i. assessed by H&E staining, scale bar 100 um. **i**) Viral titer assessed by plaque assay. The difference between genotypes was statistically not significant. n=5-9/timepoint. **j**) Flow cytometric analysis of tissue-resident (TRAMs) and infiltrating monocyte-derived alveolar macrophages (MoAMs) out of the total lung CD45+ cell population normalized to body weight during the IAV response, n=4/timepoint. **k**) Levels of inflammatory cytokines in BALF were assessed using a custom multiplex ELISA. n=3-4/timepoint. Bar graphs represent mean, error bars represent SD, differences between genotypes on each day were assessed by t-test. * p<0.05

Next, we used a clinically relevant, causal murine model of viral pneumonia to determine whether the host response to influenza A virus (IAV) infection is altered in *Vim^-/-^* mice compared to WT mice. WT and *Vim^-/-^* mice were infected with a lethal dose of IAV (Strain A/WSN/1933 (H1N1), 200 pfu) and assessed for morbidity and mortality. WT mice had a 100% mortality rate, whereas >80% of the *Vim^-/-^* mice were alive 10 days post-infection (p.i.) (**Figure 1e**). By 14 days p.i., 50% of the surviving *Vim^-/-^* mice had recovered from mild lethargy, coat ruffling, febrile shaking, and eye-watering. *Vim^-/-^* mice exhibited a lower wet-to-dry lung weight ratio and decreased total protein concentration in BALF than WT mice infected with IAV, indicating less damage to the alveolar-capillary barrier (**Figure 1f-g**). Histopathological examination of IAV-infected *Vim^-/-^* mice lungs also demonstrated less severe alveolar damage, characterized by reduced alveolar edema and less lung tissue damage than in IAV-infected WT mice (**Figure 1h**). Vimentin-deficiency did not interfere with IAV replication or clearance, as evidenced by comparable viral titers (**Figure 1i**) and similar levels of the H1N1 viral RNA molecules, such as matrix protein 2, nucleoprotein, and neuraminidase, which were detected in recruited MoAMs from WT and *Vim^-/-^* mice by RNA-seq (**Supplemental Figure 1b**). These results show that *Vim^-/-^* mice are surprisingly resilient to IAV infection with improved survival rates and decreased lung injury, independent of viral load.

The influx of immune cells into the airspace during IAV infection and the resulting production of pro-inflammatory cytokines is a major driver of lung injury. We performed a systematic flow cytometric analysis of whole-lung lysates to characterize the immune cell recruitment into the lungs (**Supplemental figure 1c**). The number of tissue-resident alveolar macrophages (TRAMs) rapidly decreased after IAV infection at a similar rate in both genotypes, with a more than 80% reduction by 6 days p.i. (**Figure 1j**). MoAMs are typically not present in the alveolar compartment in normal, healthy lung tissue at steady-state (e.g., 0 days p.i.). During IAV infection, monocytes are recruited and rapidly differentiated into MoAMs in the lung parenchyma [5, 28]. WT and *Vim^-/-^* mice infected with IAV recruited comparably increasing number of MoAMs into the lung from 2 days p.i. onward (**Figure 1j**). Similarly, neutrophil recruitment peaked on day 3 p.i. in both genotypes (**Supplemental Figure 1d**). Despite similar immune cell recruitment, *Vim^-/-^* mice had decreased levels of pro-inflammatory cytokines and chemokines, such as IL-1β, TNFα, RANTES, MIP-2, MCP-1, and Type I interferons IFNα/β, compared to WT mice following IAV infection (**Figure 1k**). In summary, wild-type mice, but not *Vim^-/-^* mice, exhibited a pro-inflammatory phenotype in response to IAV infection. As there was no difference in MoAM recruitment between genotypes, these data suggest that vimentin plays an essential role in determining the pro-inflammatory function of MoAMs.

### Three distinct temporal patterns characterize the transcriptional response of MoAMs to influenza A virus

To investigate the response of WT MoAMs to IAV infection, we performed RNA-seq on flow-sorted MoAMs to assess global gene expression profiles on 2, 3, 4, and 6 days p.i. (**Supplemental Figure 2a, 2b**). We identified 2623 differentially expressed genes over this time course and, using k-means clustering, defined three clusters that correspond temporally to three phases of the IAV response: Infiltrating (2 days p.i.), Early Inflammatory (3-4 days p.i.), and Late Inflammatory (6 days p.i.) (**Figure 2a**).

**Figure 2.**
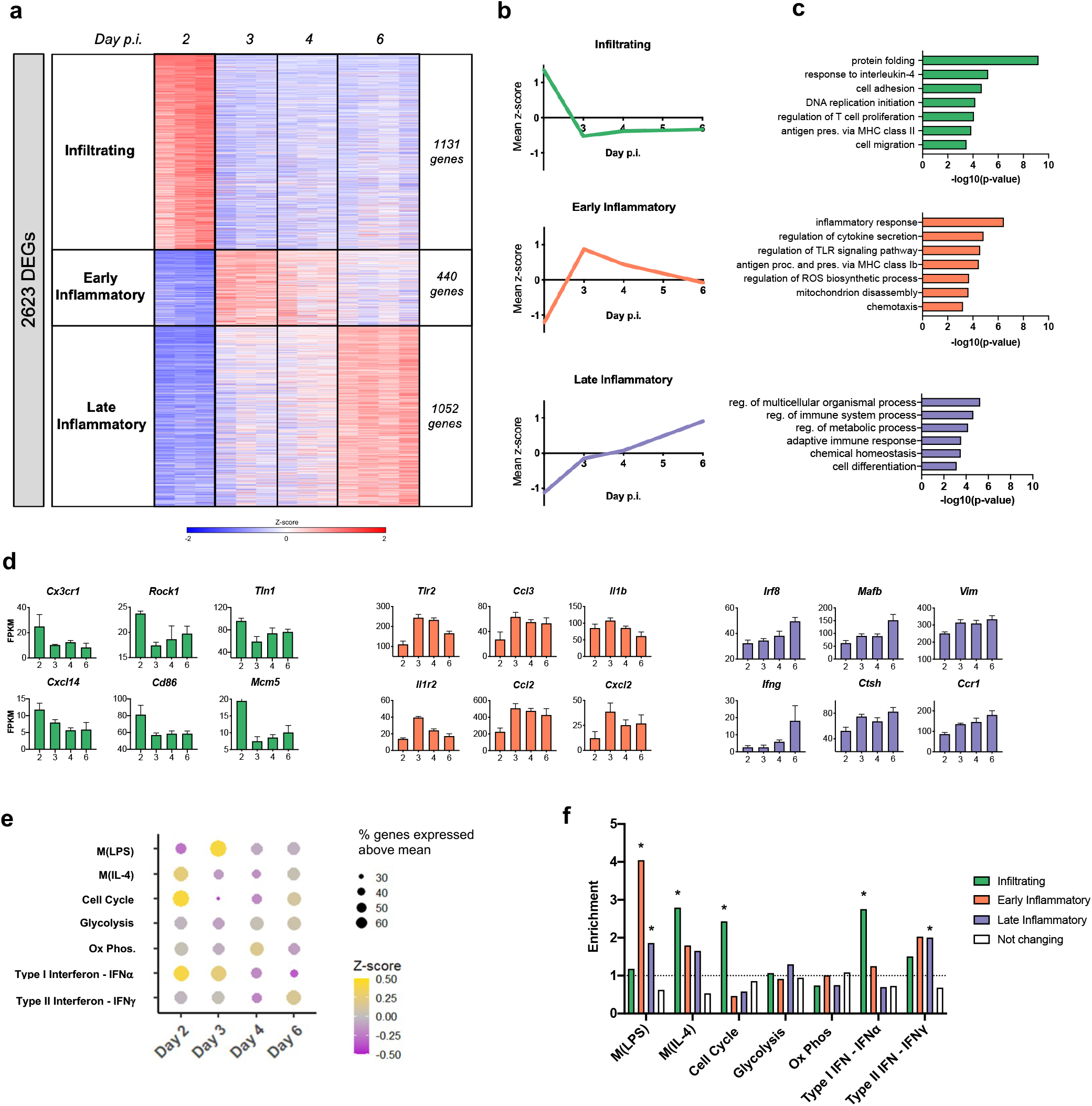
Wild-type monocyte-derived alveolar macrophages exhibit gene expression patterns associated with three phases of influenza A viral infection. **a**) Heatmap of normalized expression of 2,623 differentially expressed genes (ANOVA p-value <0.05). Z-score normalization for each row was applied to gene expression (FPKM) data. K-means clustering (k=3) identified three clusters: Infiltrating (1131 genes), Early Inflammatory (440 genes), Late Inflammatory (1052 genes). **b**) Mean z-scores calculated across genes for each cluster **c**) Gene Ontology (GO) term enrichment for each phase. **d**) Bar graphs of gene expression (FPKM) for representative genes in each phase. Error bars represent SD. (ANOVA p-value <0.05). **e**) Bubble plot representing the proportion of genes (%) within each process that had an expression above the row mean. Color gradient represents the mean z-score of expression across genes. **f**) Enrichment of gene sets in each cluster as defined in A. Asterisk represents significantly enriched overlaps (hypergeometric p-value <0.0018, FWER = 0.05). The dotted line indicates an expected enrichment score of 1. n=3-4 per group. Ox Phos = Oxidative phosphorylation.

The first cluster is characterized by genes with the highest expression in the **Infiltrating Phase** at 2 days p.i. (**Figure 2a-b, Supplemental Figure 2c**). Gene Ontology (GO) analysis finds this cluster is enriched for genes associated with *cell migration* and *cell adhesion*, such as *Cx3cr1, Rock1, Tln1, and Cxcl14* (**Figure 2c-d**), likely involved in transendothelial migration of leukocytes[29–32]. These results are consistent with the requirement for MoAMs to leave the vasculature and migrate across the endothelium to reach the alveolar space soon after IAV infection. We also observed genes in this cluster associated with the recruitment of lymphocytes, such as *Cd86*, MHC Class II gene *H2-Aa*, and *Ccl8* (**Figure 2c-d, Supplemental Figure 2d**). Furthermore, our results suggest that MoAMs may be proliferating during this phase due to the enrichment of GO processes, such as *DNA replication initiation*, and the expression of cell cycle genes, such as *Mcm5* (**Figures 2c-f**). Taken together, these data show that the primary role of MoAMs in the Infiltrating Phase of the IAV response is to enter the lung parenchyma and promote the recruitment of immune cells.

The second cluster is characterized by genes that peak in expression during the **Early Inflammatory Phase** at 3 days p.i. (**Figure 2a-b, Supplemental Figure 2c**). This cluster is enriched for genes associated with *inflammatory response* and *regulation of cytokine secretion* (**Figure 2c**) – such as *Tlr2*, *Il1b*, and *Il1r2* (**Figure 2d**). These gene expression changes correspond with the secretion of inflammatory cytokines, such as IL-1β and TNFα, beginning on Day 3 of IAV response (**Figure 1k**) and binding of viral PAMPs through pattern recognition receptors (PRRs), including TLR2 [28]. The expression of genes associated with *chemotaxis*, such as *Ccl3, Ccl2*, and *Cxcl2*, also peak during the Early Inflammatory phase (**Figure 2c-d**) and corresponds with the secretion of their respective proteins MCP-1, MIP1-a and MIP-2 measured in the BALF (**Figure 1k**). These data collectively suggest that WT MoAMs adopt a pro-inflammatory state during the Early Inflammatory Phase of the IAV response.

The third cluster is characterized by genes that display a continuous increase in expression until the **Late Inflammatory Phase** at 6 days p.i. (**Figure 2a-b, Supplemental Figure 2c**). This cluster is enriched for genes associated with the GO processes of *cell differentiation* and *adaptive immune response* – such as *Irf8*, *Mafb*, *Ifng*, *Ctsh*, *Ccr1* and, *Arg2* (**Figure 2d, Supplemental Figure 2d**). The increasing expression of *Irf8* and *Mafb* further supports the notion that recruited monocytes differentiate into monocyte-derived alveolar macrophages [33, 34]. Interestingly, the gene encoding vimentin increased in expression over time, suggesting that vimentin may play a role throughout the IAV response (**Figure 2d**). Although GO analysis implicated metabolic processes in this cluster (**Figure 2c**), and macrophage activation has been associated with cellular respiration changes[35–37], we did not observe preferential expression of genes in the glycolysis or oxidative phosphorylation pathways at any time point (**Figure 2e-f**).

Since interferon pathways are an essential element of the viral immune response, we investigated the temporal expression of Type I and Type II interferon genes. We found Type I IFNa response genes peaked in expression on day 2, which coincided with their enrichment in the Infiltrating cluster (**Figure 2e-f**). This is also consistent with Type I IFNs’ role in viral clearance. In contrast, Type II IFNy response genes did not reach their highest expression level until 6 days p.i. and were enriched in the Late Inflammatory cluster (**Figure 2d-f**). Type II IFNy has been shown to stimulate macrophages and monocytes to produce more chemoattractants and other inflammatory molecules, resulting in a cycle of persistent inflammation [1, 38]. These data support Type I interferon’s role as an early effector of the immune response, while Type II interferon is involved in the later immune response.

While GO analysis can be informative, many terms, such as *inflammatory response* and *innate immune response*, lack cell-type specificity. *In vivo* macrophages are characterized by considerable diversity and plasticity and respond to environmental cues by acquiring distinct functional phenotypes. *In vitro* macrophages can be polarized toward a pro-inflammatory (M1) phenotype or a reparative (M2) phenotype through the administration of LPS or IL-4, respectively [25, 39, 40]. To further define the generic *inflammatory response* in recruited MoAMs, we generated annotations for M_(LPS)_ and M_(IL-4)_ responses by integrating gene sets from several published studies on LPS (M_(LSP)_), and IL-4 (M_(IL-4)_) stimulated macrophages *in vitro* (see Methods)[25, 39–41]. We found that M_(IL-4)_ genes were enriched in the Infiltrating cluster, and their expression peaked on day 2 p.i. Notably, recruited WT MoAMs quickly downregulated M_(IL-4)_ genes on days 3, 4, and 6 p.i. (**Figure 2e-f**). Furthermore, we found that the M_(LPS)_ genes peaked on day 3 p.i. in WT MoAMs (**Figure 2e**) and were enriched in the Early Inflammatory cluster (**Figure 2f**). M_(LPS)_ genes were also enriched in the Late Inflammatory cluster (days 4 and 6 p.i.) but to a lesser degree. (**Figure 2f**). Taken together, our results support a critical switch in WT MoAM function as they transition from their initial transcriptional state in the Infiltrating phase to a pro-inflammatory phenotype during the Early and Late Inflammatory phases.

### *Vim^-/-^* MoAMs modulate expression of Infiltrating and Early Inflammatory, but not Late Inflammatory phase genes

To gain a better understanding of the conferred protection from IAV infection in *Vim^-/-^* mice (**Figure 1**), we performed RNA-seq on flow-sorted MoAMs from *Vim^-/-^* mice at 2, 3, 4, and 6 days p.i. (**Supplemental Figures 2a, 2b, 3a**). We compared global gene expression between genotypes each day p.i., identifying down- and upregulated genes in *Vim^-/-^* compared to WT MoAMs. The greatest differential gene expression occurred on day 3 p.i. (**Figure 3a**). To characterize processes with differential expression between genotypes, we ran GO analysis on up- and downregulated genes on each day p.i. (**Figure 3b**). Top GO processes that enrich on at least two days p.i. uncover a pattern of downregulated expression of genes involved in the inflammatory response and stress response and increased expression of genes involved in cell cycle and extracellular matrix organization in *Vim^-/-^* MoAMs throughout IAV infection (**Figure 3b**). Furthermore, using GSEA we found that gene sets associated with inflammation – including M_(LPS)_ and Type I Interferon – were significantly decreased in expression in *Vim^-/-^* MoAMs on every day p.i. (**Supplemental Figure 3b**), In contrast, *Cell Cycle* and M_(IL-4)_ associated genes had increased expression in *Vim^-/-^* MoAMs compared to WT MoAMs (**Supplemental Figure 3b**).

**Figure 3.**
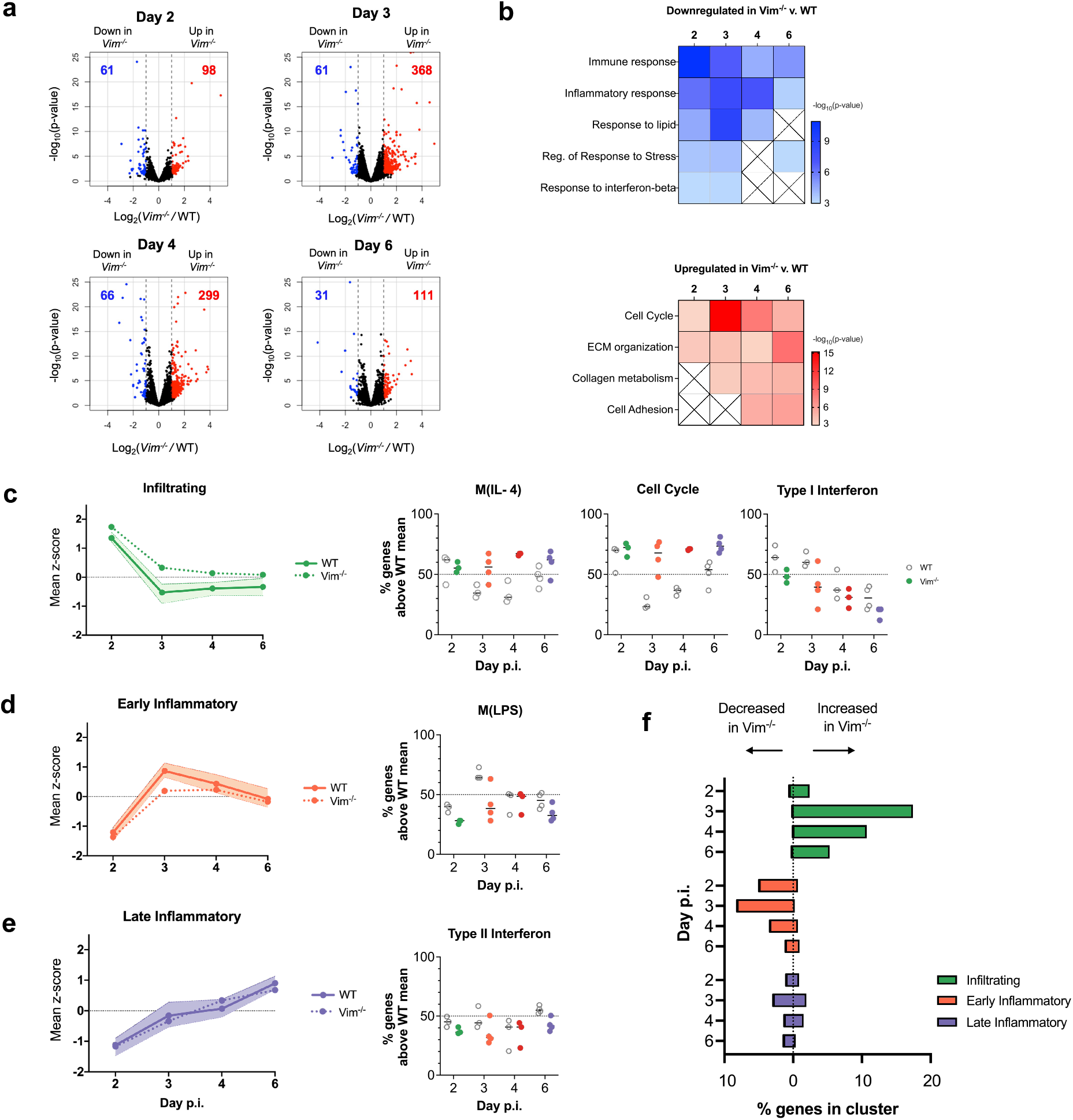
*Vim^-/-^* MoAMs modulate expression of Infiltrating and Early Inflammatory, but not Late Inflammatory phase genes. **a**) Volcano plots of differentially expressed genes (edgeR p-value<0.05, and edgeR reported log2FC>|1.0|) between *Vim^-/-^* and WT MoAMs on each day post-infection. The number of genes downregulated (blue) and upregulated (red) in *Vim^-/-^* compared to WT MoAMs are shown. **b**) GO term enrichment across days p.i. in the downregulated (blue) and upregulated (red) genes in *Vim^-/-^* compared to WT MoAMs. Scale represents −log_10_ p-value. **c-e**) Line graphs on the left represent mean adapted z-scores, calculated across genes for each cluster in WT MoAMs and *Vim^-/-^* MoAMs. Shading represents WT MoAMs inter-quartile range (IQR). Dot plots to the right represent % of genes at each timepoint expressed above their WT mean. Each dot represents a replicate. Lines represent the median **f**) % genes in each cluster that exhibit decreased or increased expression in *Vim^-/-^* MoAMs compared to WT MoAMs (n=3-4 per group). Decreased genes defined as log2FC < −0.58 (FC of 1.5); increased genes defined as log_2_FC>0.58.

Using the prior clustering of 2623 genes across the three phases of the IAV response (**Figure 2a**), we compared gene expression between MoAMs in WT and *Vim^-/-^* mice during each phase (**Figure 3c-e, Supplemental 3f-h**). In *Vim^-/-^* MoAMs, genes in the Infiltrating cluster were increased in expression while genes in the Early Inflammatory cluster tended to be decreased (**Figure 3c,d,f, Supplemental Figure 3g,h**). In contrast, genes in the Late Inflammatory cluster were less changed between genotypes (**Figure 3e, Supplemental Figure 3g,h**). In line with our previous findings, day 3 p.i. represented the most significant difference between genotypes in any cluster (**Figure 3c-f**). Indeed, day 2 and day 3 *Vim^-/-^* MoAMs were globally more similar to each other than their WT counterparts, suggesting a greater transcriptional shift occurs between day 2 to 3 p.i. in WT compared to *Vim^-/-^* MoAMs (**Supplemental Figure 3f**). These transcriptional differences between genotypes were mirrored in the data for processes associated with each phase (**Figure 2f**). M_(IL-4)_ and Cell Cycle genes, associated with the Infiltrating phase, exhibited a higher and sustained level of expression in *Vim^-/-^* MoAMs compared to WT (**Figure 3c**). These results were confirmed using GSEA (**Supplemental Figure 3b**). Type I IFN response genes were differentially expressed between genotypes, but were downregulated over time in both genotypes. In contrast, M_(LPS)_ genes associated with the Early Inflammatory phase were downregulated on 2 and 3 days p.i. in *Vim-/-* compared to WT MoAMs (**Figure 3d**), suggesting an early attenuation. Type II IFN response gene expression did not significantly differ between genotypes (**Figure 3e**). We next asked whether any of the individual gene sets correlated with each other. We observed that *M_(IL-4)_* and Cell cycle gene expression are highly correlated in both genotypes, suggesting they may be part of a more extensive network of co-regulated processes (**Supplemental Figure 3c, d**). None of the gene sets significantly differed in expression in steady-state tissue-resident alveolar macrophages, suggesting that vimentin-deficiency has a larger effect during the response to IAV-infection than in steady-state (**Supplemental Figure 3e**).

These results support an altered function of *Vim^-/-^* MoAMs in response to IAV that favors the maintenance of Infiltrating phase genes and the suppression of Early Inflammatory phase genes to confer protection from influenza-induced lung injury. Of note, these data demonstrate that the greatest difference between genotypes occurs on day 3 p.i. and this represents the key timepoint for the protective phenotype in *Vim^-/-^* mice.

### *Vim^-/-^* MoAMs fail to activate core pro-inflammatory genes in the early response to IAV

Thus far, our results suggest that the maintenance of Infiltrating phase gene expression and suppression of pro-inflammatory genes in *Vim^-/-^* MoAMs, particularly in the Early Inflammatory phase, is responsible for the resilient phenotype observed in *Vim^-/-^* mice infected with IAV (see Figure 1). To identify macrophage-specific genes involved in the Early Inflammatory Phase, we focused on the *M_(LPS)_* response associated with the increased expression on day 3 p.i. in WT mice. In MoAMs from *Vim^-/-^*mice, fewer *M_(LPS)_* genes exhibited peak expression on day 3 p.i. (**Figure 4a**). Moreover, *M_(LPS)_* genes were enriched among those genes exhibiting the greatest expression decrease in *Vim^-/-^* MoAMs compared with WT during the course of IAV response (**Supplemental Figure 3b**). Of the *M_(LPS)_* genes most decreased on at least one day, we find that the majority failed to intensify expression on day 3 p.i. in *Vim^-/-^* MoAM compared with WT MoAM (**Figure 4b-c, Supplemental Figure 4a**). This core set of pro-inflammatory genes includes interferon-stimulated genes (ISGs), such as *Ifit1*, *Irgm1*, and *Rsad2*, genes associated with chemokine pathways, including *Ccrl2* and *Cxcl2*, and genes associated with pathogen sensing, *Fcgr1, and Tlr2* (**Figure 4b-c**). Complementary to the decrease in *M_(LPS)_* gene expression, *M_(IL-4)_* genes, such as *Cd36*, *Cd11c*, and *Cd83*, previously associated with the Infiltrating phase, exhibited higher expression on all days post IAV infection in *Vim^-/-^* MoAMs compared with WT MoAMs (**Supplemental Figures 3b, c, 4b**). Collectively, these data suggest *Vim^-/-^* MoAMs fail to activate core pro-inflammatory genes associated with the Early Inflammatory phase in response to IAV.

**Figure 4.**
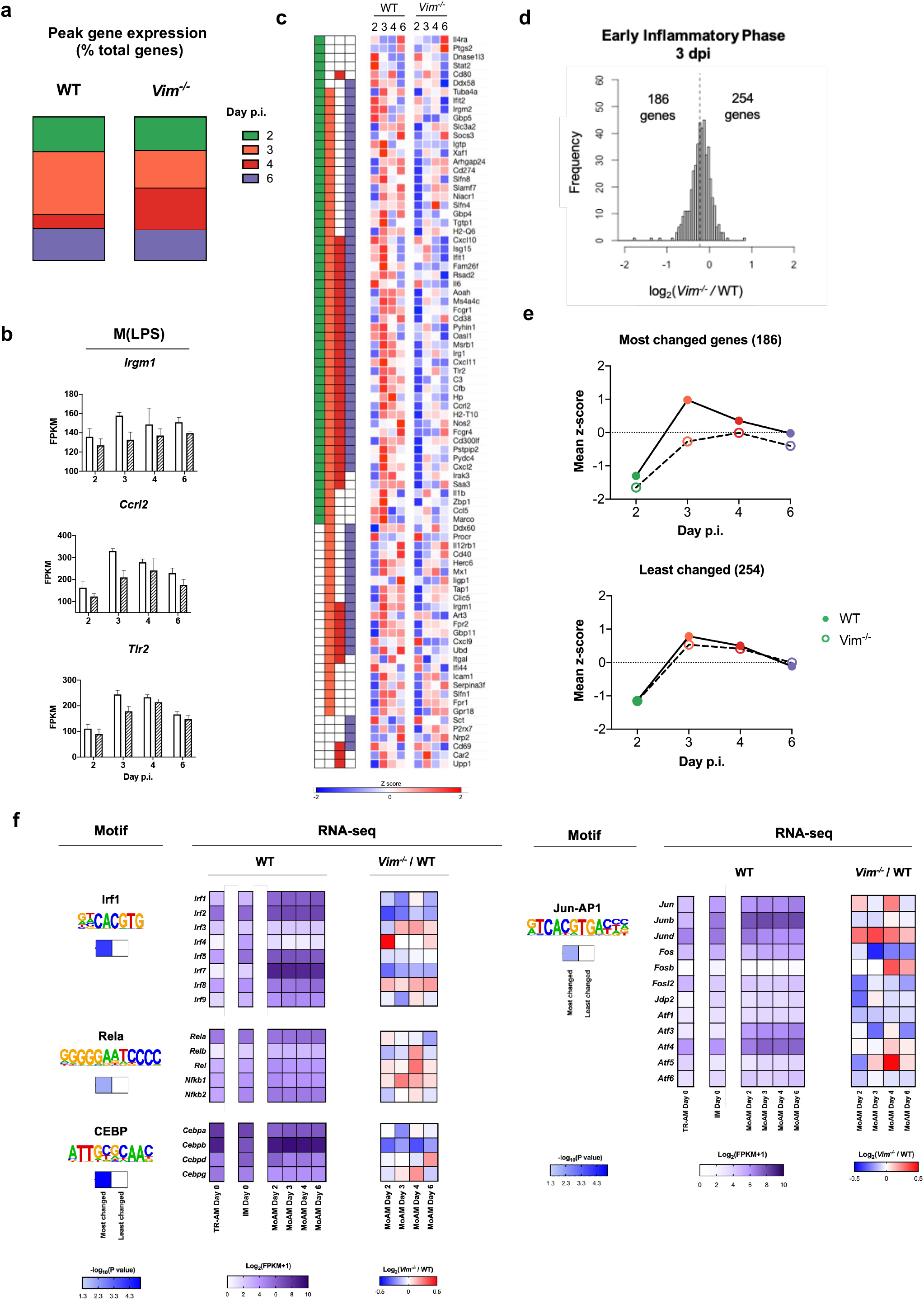
Protective phenotype of *Vim^-/-^mice* is associated with the failure of *Vim^-/-^MoAMs* to activate core pro-inflammatory genes in the early response to IAV. **a**) Percent (%) M_(LPS)_ genes peaking in expression on each day post-infection in WT and *Vim^-/-^* MoAMs **b**) Bar graphs of gene expression (FPKM) for representative M_(LPS)_ genes present in the leading edge of at least 1 day from GSEA. Error bars represent SD. **c**) Heatmap of normalized expression of 84 M_(LPS)_ genes present in the GSEA leading edge on at least 1 day (indicated by a color-coded bar on the left). Adapted Z-score normalization using WT mean and standard deviation for each row were applied to gene expression (FPKM) data. **d**) Histogram representing the distribution of log2FC between *Vim^-/-^* and WT MoAMs on 3 dpi of the Early Inflammatory phase (440 genes). The dashed line represents the mean of the distribution. The number of genes less than (186) and greater than (254) the mean are shown. Genes with log2FC between *Vim^-/-^* and WT MoAMs less than the mean represent those genes that are most changed between *Vim^-/-^* and WT MoAMs. **e**) Mean adapted z-scores were calculated across genes for most (186) and least (254) changed genes on 3 dpi in the Early Inflammatory Phase in WT and *Vim^-/-^* MoAMs. **f**) HOMER known motif analysis using genomic background of most changed genes on 3 dpi in the Early Inflammatory Phase (186 genes). Motif logos are shown below motif names. −logiopvalues of enrichment in most or least changed gene set is shown. Log_2_(FPKM+1) average gene expression of transcription factor family members are shown for WT TRAMs and WT IMs at steady-state (0 dpi) and 2-6 dpi for WT MoAMs. log_2_FC between *Vim^-/-^* and WT MoAM gene expression of transcription factor family members on each dpi. (n=3-4)

Prior work has demonstrated that macrophage activation is largely defined by the activity of cell-type-specific transcription factors (TF) that poise regulatory elements for activation by stimulus-specific TFs [42–44]. To better characterize the impact on gene regulation observed in *Vim^-/-^* MoAMs, we searched for TF binding sites enriched in the promoters of vimentin-dependent genes associated with the Early Inflammatory phase. First, we classified the 440 Early Inflammatory genes into those “most changed” and “least changed” in *Vim^-/-^* MoAMs based on their fold-change compared with WT MoAMs (**Figure 4d-e, Supplemental Figure 4c**). Next, we performed motif-finding on each gene set using a known sequence motifs database (see Methods). We found that TF families previously implicated in an inflammatory response, such as *Jun-AP1* and *Rela*, were significantly enriched among the most changed but not the least changed genes (**Figure 4f**). Some Irf family motifs were differentially enriched between the two gene sets, while others were shared (**Figure 4f**). As supported by induction of gene expression in IAV and the decrease observed in *Vim^-/-^*, those associated with ISGs (*Irf1/7/9*) were more likely to be affected by vimentin-deficiency (**Figure 4f**). At the same time, *Irf8*, which is more generally involved in macrophage function, exhibited similar enrichment in both groups (**Supplemental Figure 4e**). Furthermore, the CEBP motif was associated with the most changed genes in *Vim^-/-^* MoAMs, and *Cebpb* was expressed highly at both baseline and throughout IAV infection (**Figure 4f**). This observation is consistent with *Cebpb*, a known macrophage TF that can also act in combination with AP1 factors (**Figure 4f**), specifying poised regulatory elements in macrophages that activate genes on stimulation[45, 46]. On the other hand, ETS was the main TF family with motifs enriched in the least changed but not the most changed genes (**Supplemental Figure 4d**). This family marginally includes PU.1, a well-known myeloid lineage factor, but PU.1 is unlikely to have a specific role in IAV response (**Supplemental Figure 4e**). Instead, the expression of *Elf1* and *Elf4* suggests that they may have an interferon-independent antiviral role in influenza as previously reported [47], that is not impacted by vimentin-deficiency. In summary, vimentin-deficiency impedes the stimulus-dependent regulation of pro-inflammatory genes in MoAMs during the Early Inflammatory phase of IAV infection. Although this effect leads to subtle changes in overall gene expression, it appears sufficient to alter the MoAM function.

### Vimentin-deficient MoAMs are sufficient to confer protection from IAV-induced mortality

We examined whether the loss of vimentin *in vitro* paralleled our *in vivo* observations of suppressed pro-inflammatory gene expression (**Figure 3,4**). We polarized bone marrow-derived macrophages (BMDMs) from WT and *Vim^-/-^* mice toward a pro-inflammatory (M1/M_(LPS)_) phenotype or a reparative (M2/M_(IL-4)_) phenotype through the administration of LPS or IL-4, respectively (**Figure 5a**). In line with our *in vivo* observations, WT M1/M_(LPS)_ macrophages had increased expression of pro-inflammatory genes, such as *Tnf* and *Il1b*. The loss of vimentin suppressed the expression of M1 pro-inflammatory genes (**Figure 5b**) and increased the expression of M2/M_(IL-4)_ reparative genes (*Mrc1, Arg1*) compared to WT macrophages (**Figure 5b**). Collectively, these data suggest that vimentin plays a critical role in defining the macrophage pro-inflammatory phenotype.

**Figure 5.**
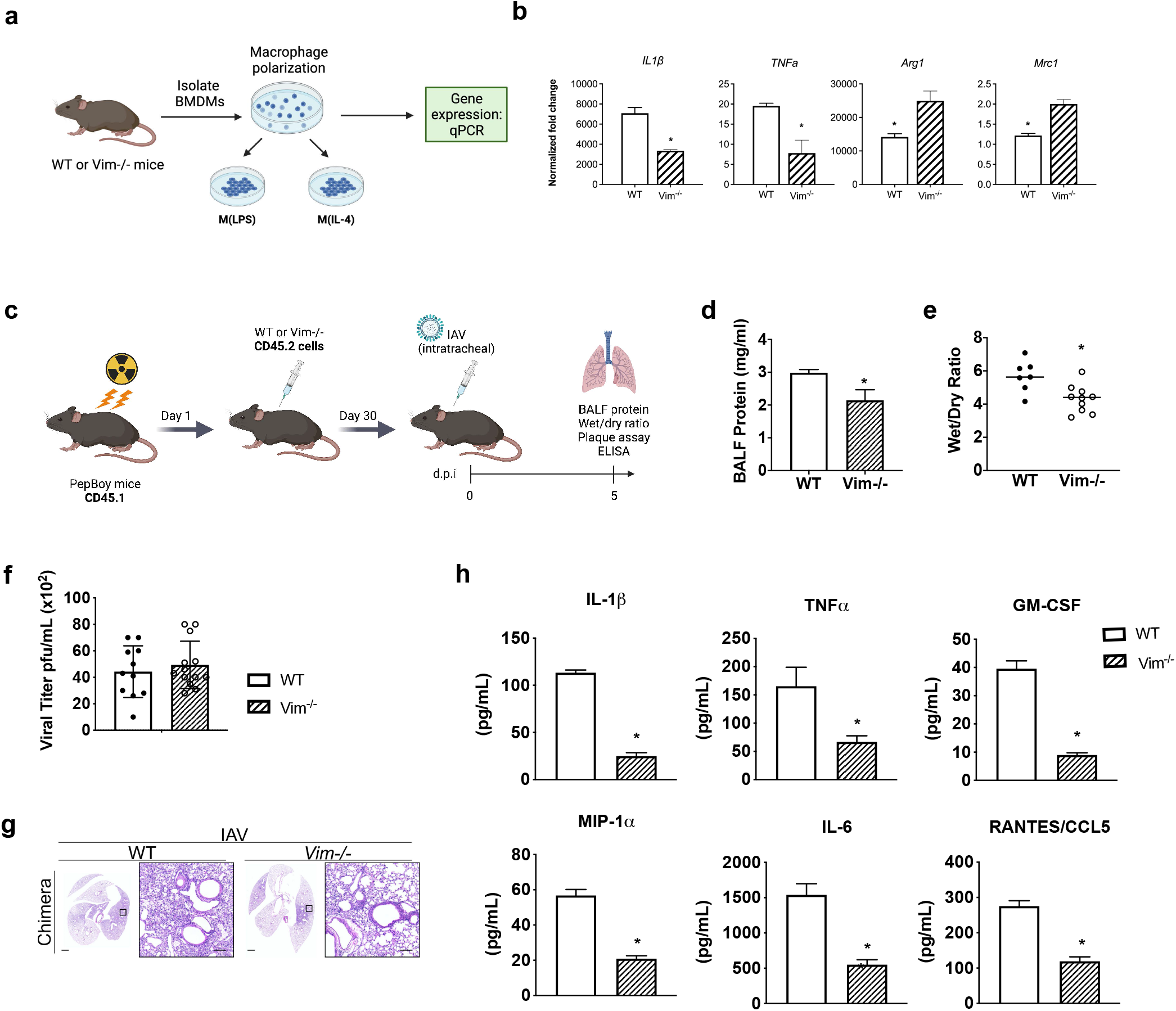
Vimentin-deficiency in the MoAM compartment is sufficient to protect from influenza A virus-induced lung damage and mortality. **a**) Schematic of experimental design. BMDM were isolated from either WT or Vim-/- mice. BMDM were treated with LPS or IL-4 to polarize macrophages towards an M_(LPS)_ or M_(IL-4)_ state, respectively. Gene expression was assessed by qPCR **b**) Expression of genes characterizing a pro-inflammatory and reparative macrophage state in WT and *Vim^-/-^* macrophages. n = 2, in duplicate. Bar graphs represent mean relative fold change, error bars represent SEM. Differences between genotypes were assessed by t-test. **c**) Schematic of the experimental design. Recipient mice (CD45.1) were irradiated and reconstituted with bone marrow cells from WT or *Vim^-/-^* donor mice (CD45.2). After a 30-day recovery period, chimeric mice were intratracheally (i.t.) infected with a lethal dose of IAV at 5 days p.i. and subjected to analyses. **d**) Bronchoalveolar lavage fluid (BALF) protein levels measured by Bradford assay, n=3-4 **e**) Lung wet-to-dry ratio measured at day 5 p.i. n=7-10. **f**) Viral titer assessed by plaque assay. n=11-14. **g**) Representative images of murine lung architecture at day 5 p.i. assessed by H&E staining, scale bar 100 um. **h**) Levels of inflammatory cytokines in BALF assessed using a custom multiplex ELISA. n=4-9. Bar graphs represent mean, error bars represent SEM. Differences between genotypes were assessed by t-test. * p<0.05.

To determine if vimentin plays a causal role in defining the pro-inflammatory function of recruited MoAMs we used isochronic adoptive MoAM transfer experiments and molecular profiling in IAV-infected mice. Bone marrow derived cells from WT or *Vim^-/-^* donor mice (CD45.2) were transferred into irradiated WT recipient mice (CD45.1), allowing for the specific reconstitution of MoAMs with a WT or *Vim^-/-^* genotype (**Figure 5c**). All mice showed reconstitution of donor hematopoietic cells in peripheral blood above 85% (**Supplementary Figure 5a**). Following confirmed engraftment and recovery, mice were infected with a lethal dose of IAV (**Figure 5c**). We found that IAV-infected WT mice who received *Vim^-/-^* donor bone marrow exhibited significantly lower total protein concentration in the BALF and lower wet-to-dry lung weight ratio than mice that received WT donor bone marrow, indicating less severe alveolar damage (**Figure 5d-g**). Mice showed no difference in IAV replication or clearance regardless of donor genotype (**Figure 5f**). Histopathological examination of IAV-infected WT mice who received *Vim^-/-^* donor bone marrow demonstrated less severe alveolar damage than those who received WT bone marrow (**Figure 5g**). As in *Vim^-/-^* mice (**Figure 1**), WT mice with *Vim^-/-^* MoAMs exhibited lower levels of pro-inflammatory cytokines and chemokines (**Figure 5h**). Taken together, these data support the notion that vimentin plays a causal role in determining the pro-inflammatory function of recruited MoAM and drives IAV-induced lung injury.

## Discussion

Monocytes are recruited to the lung during injury, where they differentiate into monocyte-derived alveolar macrophages (MoAMs) in response to cues provided by the injured microenvironment. We used a clinically relevant murine model of severe viral pneumonia to identify monocyte-derived alveolar macrophage-specific gene expression patterns associated with three distinct phases of IAV infection: Infiltrating, Early Inflammatory, and Late Inflammatory. WT MoAMs began expressing pro-inflammatory genes and secreting tissue-damaging pro-inflammatory cytokines in the Early Inflammatory phase, resulting in increased morbidity and mortality in influenza-infected wild-type mice. Conversely, upon entering the IAV-infected alveolar microenvironment, *Vim^-/-^* MoAMs failed to upregulate pro-inflammatory genes and suppressed the secretion of pro-inflammatory cytokines conferring protection to the host. By leveraging the *Vim^-/-^* mouse’s protective phenotype, we investigated the transcriptional response following IAV infection and identified a core set of pro-inflammatory genes that drive lung injury and mortality. Although prior studies have examined macrophage gene expression at steady-state or single time points following IAV infection [48, 49], by analyzing gene expression over a time course post-IAV infection, our results identified critical temporal dynamics that shape the function of recruited monocyte-derived alveolar macrophages in response to respiratory infections.

During IAV-infection, infected cells, and cells dying from necroptosis or pyroptosis, release PAMPs and DAMPs, which activate signaling pathways to recruit additional effector cells, including MoAMs. We reason that as monocytes extravasate and infiltrate the lung parenchyma, the differentiation of WT MoAMs toward a pro-inflammatory phenotype involves activation of specific transcriptional programs in response to the IAV-infected lung microenvironment. Understanding how MoAMs communicate with resident cell populations to drive persistent inflammation (e.g., WT MoAMs) or promote reparative processes (e.g., *Vim^-/-^* MoAMs) represents an active area of investigation. During homeostasis, alveolar epithelial cells interact with tissue-resident macrophages via receptor/ligand pairs CD200/CD200R, signal regulatory protein-α (SIRPα)/CD47, and CSF2R/GM-CSF and immune/epithelial E-cadherin interactions [50, 51]. It follows that similar interactions occur between alveolar epithelial cells and recruited MoAMs in the infected lung. These interactions, or the disruption of these interactions, may activate specific transcriptional programs contributing to the pro-inflammatory phenotype we observed in WT MoAMs. Investigating receptor/ligand interactions and the secretion of molecules is limited by their short half-life and the requirement for mass spectroscopy for their detection. Importantly, little information is known about specific receptor/ligand interactions in recruited MoAMs during lung injury. We demonstrated that WT and *Vim^-/-^* MoAMs encountered similar environmental signals associated with IAV infection (e.g., equivalent viral load, similar immune cell recruitment – see Figure 1). Notably, *Vim^-/-^*MoAMs suppress pro-inflammatory genes and cytokines and maintain an increased expression of reparative M_(IL-4)_ genes throughout IAV infection. We hypothesize that the protective phenotype displayed by recruited *Vim^-/-^* MoAMs may be the result of disrupted or altered receptor/ligand interactions in the IAV-infected lung microenvironment. It will be important to investigate how the temporal transcriptional changes observed in our study impact the ability of recruited MoAMs to interact with and respond to other cell types in the lung microenvironment.

*In vitro* studies have established vimentin’s role in essential cellular processes, such as cell motility, cell signaling, and protein complex scaffolding [52–55]. Retinoic acid-inducible gene I, a cytosolic PRR that detects viral RNA and mediates type I interferon expression, leads to increased vimentin expression[56]. Our data support this previous observation, as type, I interferon genes peaked on day 2 p.i., preceding the increase in vimentin gene expression in WT MoAMs. We confirmed previous reports that *vimentin-deficient* mice fail to secrete mature IL-1b as assembly and activation of the NLRP3 inflammasome requires vimentin [18, 19]. Complementary to our findings, vimentin’s interaction with NOD2, a cytoplasmic PRR, activated NF-κB and induced a pro-inflammatory immune response in an inflammatory bowel disease model [20]. Of note, the suppressed pro-inflammatory response observed in *Vim^-/-^* MoAMs does not appear to result from differing macrophage maturation. In fact, genes involved in the transition from monocyte to macrophage-like state, such as *Mafb*, were not differentially expressed between genotypes. Collectively, this suggests that vimentin mediates the innate immune response(s) by regulating the expression of pro-inflammatory genes.

Our findings demonstrate that depleting vimentin in MoAMs modulates the pro-inflammatory response and protects the host, providing a rationale that vimentin could serve as a potential therapeutic target during viral infection. Our group and others have shown that vimentin can be therapeutically targeted in non-small cell lung cancer and other diseases such as idiopathic pulmonary fibrosis [57, 58]. Molecules such as withaferin A, FiVe1, and simvastatin were used to perturb the expression and/or assembly of vimentin filaments and led to the downregulation of vimentin was accompanied by decreased cancer progression, decreased fibrosis and intermediate filament disorganization, and cell death [58–61]. Collectively, these and our studies serve as a proof-of-principle to support the development of pharmacological agents to target vimentin selectively. These tools may be clinically beneficial in modulating the hyperinflammatory microenvironment of the virus-infected lung.

Our study has several limitations. We used a well-established bone-marrow chimera mouse model to demonstrate that *Vim^-/-^* MoAMs failed to upregulate pro-inflammatory genes and secrete pro-inflammatory cytokines, which was sufficient to confer protection from IAV-induced lung injury and mortality. Additionally, we showed that *Vim^-/-^* BMDMs expressed higher levels of M2/M_(IL-4)_ markers and lower levels of M1/M_(LPS)_ markers, which appears to reflect the macrophage plasticity we observed in our murine model of viral pneumonia. We caution the reader that while the framework of M1/M2 polarization has provided a useful system to study macrophages in vitro, it is unlikely that this occurs in tissue. This was shown by Xue *et al*., who found that the M1/M2 paradigm failed to describe the transcriptome of human monocyte-derived and alveolar macrophages stimulated with LPS/IFN-γ or IL-4/IL-13 in the presence of factors that recapitulate different tissue or disease microenvironments [62]. Future studies could be improved by using a lineage mark to track the recruitment and phenotype changes associated with recruited MoAMs *in vivo*. By generating a *Vim^fl/fl^Cx3cr1^CreERT2^zsGreen* transgenic mouse, we could selectively delete vimentin and track the recruitment of *GFP^+^Vim^-/-^* alveolar macrophages during IAV infection. Regrettably, the vimentin-floxed mouse is not yet available. Second, our RNA-seq data identified distinct transcriptional phases of the IAV response in MoAMs; however, it is unclear whether different MoAM subpopulations enter the lung at different time points from a single compartment. For example, MoAMs may differentiate directly from circulating monocytes that extravasate from the pulmonary capillaries into the alveolar space. Alternatively, MoAMs may stem from interstitial macrophages already residing in the lung at steady-state. Using a *Vim^fl/fl^Cx3cr1^CreERT2^zsGreen* mouse, pulsed lineage tracing would address these important questions.

In summary, our results define the temporal, transcriptional dynamics in recruited alveolar macrophages that shape the response to respiratory infection and demonstrate that vimentin plays an essential role in modifying macrophage phenotype and function. Moreover, we defined a core set of pro-inflammatory genes whose upregulation during the early response to IAV drives lung injury and mortality in wild-type, but not *Vim^-/-^* mice. We show that vimentin-deficient recruited alveolar macrophages exhibit a dampened inflammatory response in a cell-autonomous manner, sufficient to confer protection from IAV-induced mortality. Our data demonstrate that vimentin plays a causal role in shaping the pro-inflammatory response in recruited MoAMs, promoting virus-induced lung injury following IAV infection.

## Materials & Methods

### In silico single-cell RNA-seq data analysis

We queried a single-cell RNA-seq transcriptomic data set derived from bronchoalveolar lavage samples from 10 patients with severe SARS-CoV-2 viral pneumonia[1]. We downloaded normalized counts tables and visualized the pre-determined cell population clusters using Scanpy [63]. 40,000 single cells representing the four monocyte-derived alveolar macrophages (MoAMs) populations were visualized as a UMAP. Mean expression data of individual genes per cell population cluster were visualized as dot plots in Scanpy.

### Mice

All experiments were performed using wild-type 129S or vimentin-null (*Vim^-/-^*) 129S/V mice, age 8-12 weeks. Wild-type mice were purchased from Taconic, *Vim^-/-^* mice were a gift from Albee Messing (University of Wisconsin, Madison, WI). All animals were kept under pathogen-free conditions, and all experiments were approved in accordance with Northwestern University’s Institutional Animal Care and Use Committee (IACUC) guidelines.

### Bone marrow chimeras

8-12 week old recipient mice (Ptprc^a^/Pep-Boy, CD45.1) were lethally irradiated with a single dose of 1,000 cGy γ-radiation using a Cs-137–based Gammacell-40 irradiator (Nordion). Within 24 hours, fresh bone marrow cells (2 × 10^6^) from either WT donor mice or *Vim^-/-^* donor mice (CD45.2) were injected via retro-orbital injection. Mice were housed together in individually ventilated cages and were given *ad libitum* access to food and water supplemented with Trimetoprim/Sulfamethoxazole (TMP/SMX 40/8 mg formula, Hi-Tech Pharmacal) for 4 weeks and then switched to standard housing regimen. Reconstituted mice were used for influenza infection experiments ~4 weeks after transplantation. Engraftment was assessed after two weeks, using flow cytometry to measure the percentage of CD45.1+ and CD45.2+ cells in peripheral blood collected from the facial vein. All mice used for experiments showed reconstitution of the donor (CD45.2) circulating leukocytes in peripheral blood above 85%.

### Bone marrow derived macrophage (BMDM) polarization and RT-PCR

BMDMs were isolated from wild-type 129S or vimentin-null (*Vim^-/-^*) 129S/V mice, age 8-12 weeks, and treated with 100 ng/mL LPS or 20 ng/mL IL-4 to induce M_(LPS)_ and M_(IL-4)_ polarization, respectively. Cells were lysed after 24 hours and RNA was extracted using RNeasy Mini kit (Qiagen) according to manufacturer’s protocol. RNA concentration was assessed using a NanoDrop platform (Thermo Fisher). RT-PCR was run to assess gene expression changes (BioRad CFX).

### Virus

H1N1 Influenza virus strain A/WSN/1933 (WSN) was propagated in fertile chicken eggs (Sunnyside Hatchery) as previously described[4, 5]. Viral titers were determined by measuring plaque-forming units (pfu) in Madin-Darby canine kidney epithelial cells (ATCC).

### Plaque Assay

Confluent monolayers of Madin-Darby canine kidney epithelial cells were infected with stock WSN virus in 1% BSA DMEM for 2h at 37°C. Plates were washed with PBS and an overlay of 50% 2x replacement media (2x DMEM, 0.12M NaHCO3, 2% penicillin-streptomycin, and 1% HEPES), 50% Avicel (2.35%), and N-acetyl trypsin (1.5μg/mL) remained on the cells for 72h at 37°C. Plates were washed with PBS and fixed with 0.2% PFA before staining the monolayers with naphthalene blue-black stain.

### In vivo influenza virus infection

8-12 week old WT and *Vim^-/-^* 129S/V male mice were anesthetized using isoflurane and intratracheally (i.t.) infected with 200 pfu WSN. Mice were monitored daily, and weight was recorded every 24 hours.

### Histology

Mice were euthanized, after which a 20-gauge angiocath was sutured into the trachea. Heart and lungs were removed *en bloc*, and lungs were inflated with 0.8mL of 4% paraformaldehyde (PFA) at a pressure not exceeding 16 cm H_2_O. Tissue was fixed and embedded in paraffin. 5 μm sections were stained with H&E by the Mouse Histology Phenotyping Laboratory (Northwestern University, Chicago, IL). Tissue sections were visualized using TissueGnostics, a Tissue/Cell High Throughput Imaging Analysis Systems (Vienna, Austria) and captured using TissueFAXS software (TissueGnostics, Los Angeles, CA) at the Northwestern University Cell Imaging Facility (Northwestern University, Chicago, IL).

### Bronchoalveolar Lavage

A 20-gauge angiocath was ligated into the trachea, and 1mL of sterile PBS was instilled into the lungs, then removed through the angiocath. This process was repeated three times. Bronchoalveolar lavage fluid (BALF) was centrifuged at 500 x *g* for 10 min Protein levels in the supernatant were measured by Bradford assay (Bio-Rad), and cytokine levels were measured using a custom multiplex ELISA (ThermoFisher Scientific).

### Wet-to-dry weight ratio

Mice were euthanized, and lungs were surgically removed *en bloc*. The left lung was ligated, excised, and weighed in a tared container. The left lung was dried at 70°C in a Speed-Vac SC100 evaporator (Thermo Scientific, Waltham, MA) until a constant weight was obtained, and the wet-to-dry weight ratio was calculated.

### Flow cytometry analysis of lung cell populations

Mice were euthanized, and lungs were perfused with 10mL HBSS with Ca^2+^ and Mg^2+^. Lung loves were removed and inflated with enzyme solution (5mL of 0.2 mg/mL DNase I and 2 mg/mL collagenase D in HBSS with Ca^2+^ and Mg^2+^) using a 30-gauge needle. According to the manufacturer’s instructions, tissue was minced and processed in gentleMACS dissociator tubes (Miltenyi Biotec). Processed lungs were passed through a 40μm cell strained, and RBCs were lysed using BD PharmLyse (BD Biosciences, San Jose, CA). The remaining cells were counted using a Countess cell counter (Invitrogen, Grand Island, NY). CD45 microbeads were added, and cells were eluted according to the Miltenyi manufacturer’s instructions. Cells were stained with viability dye Aqua (Invitrogen), and non-specific antibody (Ab) binding was inhibited by adding Fc Block (553142, clone 2.4G2; BD Pharmingen). Blocked cells were stained with a mixture of fluorochrome-conjugated Abs (see **Online Supplemental Table 1** for list of Abs). Data were acquired on a BD LSR II flow cytometer using BD FACSDiva software (BD Biosciences), and data analyses were performed using FlowJo software (Tree Star, Ashland, OR). Cell populations were identified using a sequential gating strategy (**Supplemental Figure 1b**). Cell number was normalized to body weight.

### Macrophage harvesting and flow sort

Mice were euthanized and perfused with 5 mL sterile PBS by cardiac perfusion through the right ventricle. A 20-gauge angiocath was ligated into the trachea, and the lung and trachea were removed *en bloc*. Lungs were instilled with 1 mL dispase (Corning) through the angiocath, and a second suture was tied around the trachea to avoid leakage. Lungs were placed in 50 mL tubes (Falcon) containing 750 μL cold dispase and rocked for 45 min at RT. Trachea and angiocath were removed, and lungs were minced using curved-edge scissors. Digested tissue was poured into 50 mL tubes containing 7 mL DMEM (Corning) and 14 μL DNase I (50 mg/mL, Sigma). Tubes were rocked for 10 minutes at RT. Using serological pipettes, tissue was gently further disrupted and subsequently filtered through 70 μm cell strainers and 40 μm strainers (Corning). Tubes were rinsed with DMEM + 5% FBS and filtered through the same strainers. Cell suspensions were centrifuged for 10 min at 1300 rpm, 4°C and pellets were resuspended in 350 μL MACS buffer (Milteny). 100 μL CD11c magnetic microbeads (Milteny) were added, and cells were eluted according to the Miltenyi manufacturer’s instructions. Cells were stained for CD11b (Biolegend), Siglec F (BD), Ly6C (eBioscience), CD45 (eBioscience), MHCII (Biolegend), CD64 (Biolegend), Ly6G (BD), and NK1.1(BD). Viability dye (eFluor506, eBioscience) was used to sort living cells only (**Online Supplemental Table 1**). Using sequential gating (**Supplemental Figure 2b**), recruited alveolar macrophages (MoAMs) and tissue-resident alveolar macrophages (TRAMs) were flow-sorted using a BD FACSaria III 5-laser cell sorter and 100 μm nozzle (BD Biosciences). Of note, MoAMs were collected in the same gate as interstitial macrophages (Ly6C-, CD64+); however, by 2 days p.i., interstitial macrophages were no longer present in this gate (see **Online Supplemental Figure 2**). Flow-sorted cells were collected into 1.5 mL Eppendorf tubes containing 200 μL MACS buffer and centrifuged at 5,000 rpm for 5 min at RT. Pellets were resuspended and lysed in RLT lysis buffer (Qiagen) and stored at −80°C until all time points were collected.

### RNA sequencing (RNA-seq)

Samples collected from infected mice that at flow sort appeared uninfected (had no infiltrating MoAM) were excluded from RNA isolation. RNA was isolated in all samples at the same time to minimize the batch effect. All libraries for a given cell type were prepared as a single batch for sequencing. Total RNA was isolated using RNeasy Plus Mini kits (Qiagen). RNA quality (RIN) and quantity were assessed using a TapeStation 4200 (Agilent Technologies). RIN scores ranged from 7.2 to 9.4, with an average of 8.6. Using poly(A) mRNA magnetic enrichment kits from NEBNext (New England Biolabs), mRNA was isolated. cDNA libraries were prepared using NEBNext Ultra DNA library kits (New England BioLabs) for Illumina. Libraries were sequenced using high-output 75-cycle single-end kits on an Illumina NextSeq 500 sequencer to an average depth of 4.1 million reads per sample. Reads were demultiplexed (bcl2fastq), trimmed, and aligned to reference genome mm10 using TopHat2[64]. Aligned reads were mapped to genes using HTSeq with an Ensembl annotation (version 78)[65].

### RNA-seq analysis

For k-means clustering, we used gene counts normalized to FPKM (Fragments per Kilobase of transcript per Million mapped reads). Low counts were removed, with a cut-off set as row sum >27 (# of samples used for analysis), yielding a gene counts table of 9,169 expressed genes. One-way ANOVA with Benjamini-Hochberg (BH) FDR correction was used to define differentially expressed genes over time. Genes with an FDR <0.05 were considered differentially expressed, resulting in 2,623 DEGs. Clustering was performed on the 2,623 DEGs using k-means clustering (GENE-E, https://software.broadinstitute.org/GENE-E/index.html).

For pairwise comparison, low counts were trimmed if CPM <2 in all replicates in the smallest group of the comparison. EdgeR was used to define pairwise differentially expressed genes between WT and *Vim^-/-^* MoAMs[66, 67]. Principal Component analysis (PCA) plots were generated using *pca* function in R. Pearson correlation matrix, and heatmaps were generated using GENE-E. Gene ontology (GO) analysis was run using Gorilla[68, 69]. Expressed genes, filtered for low counts (9,169) were used as background. Gene set enrichment analysis (GSEA) was run using the weighted pre-ranked setting based on the log2FC reported by edgeR [70, 71]. Gene sets associated with specific processes were downloaded from KEGG (Cell Cycle) or Hallmark pathways (Glycolysis, Oxidative Phosphorylation, IFN alpha (Type I interferon), IFN_γ_ (Type II Interferon) except for the custom gene sets described below. The 85 overlapping genes between the IFNa and IFN_γ_ Hallmark pathways were excluded from the IFN_γ_ gene set. Overlaps between gene lists were visualized using Venny (https://bioinfogp.cnb.csic.es/tools/venny/index.html).

### Custom gene set generation

Custom gene sets for LPS/IFN_γ_ and IL-4 stimulated were generated by querying primary published data available in relevant experimental models. The final curated gene sets, M(LPS/IFN_γ_) and M_(IL-4)_, were derived from four primary publications [25, 39–41]. All four publications used mouse bone-marrow-derived macrophages (BMDM) and stimulated them *in vitro* with either LPS+IFN_γ_or IL-4. Methodologies used included LC-ESI-MS/MS mass spectrometry, microarray, and RNA-seq. A summary of the experimental approach, methods, and results for all publications can be found in **Online Supplemental Table 2**. We included all genes that were upregulated in at least one data set and could be converted into a unique Ensembl ID. Three genes were found in both M_(LPS/IFN_γ_)_ and M_(IL-4)_ and were removed to avoid any overlap between gene sets. These trimming steps resulted in two annotated gene sets of 110 LPS/IFN_γ_ stimulated genes and 101 IL-4 stimulated genes (**Online Supplemental Table 2**).

### Motif finding

We generated two gene sets of the 186 most and 254 least changed genes in *Vim^-/-^* MoAM by splitting the 440 genes of the Early Inflammatory cluster (Figure 2) based on whether their log_2_(*Vim^-/-^/WT*) expression on day 3 p.i. was above or below the mean of −0.23. (**Figure 6d,e**). To identify motifs that were enriched in either of these gene sets, we ran findmotifs.pl from the HOMER package with default parameters and the mouse mm10 genome as background [45]. We identified significantly enriched motifs as those with p-value < 0.05 in the known motif results.

### Viral reads

To identify viral reads within libraries, unmapped reads were aligned to the H1N1 reference genome. Reads for the H1N1 matrix protein 2 (M2), nucleoprotein (NP), and neuraminidase (NA) were normalized by sample to the number of uniquely aligned (mm10) reads.

### Statistics

Analyses for cytokine levels, BALF protein, viral titer, and wet-to-dry ratio were performed using GraphPad Prism (GraphPad Software). P values were calculated using a two-tailed unpaired t-test assuming equal variance.

## Acknowledgments

Histology services were provided by the Northwestern University Mouse Histology and Phenotyping Laboratory, which is supported by NCI P30-CA060553 awarded to the Robert H Lurie Comprehensive Cancer Center. Flow Cytometry was supported by the Northwestern University RHLCCC Flow Cytometry Facility and a Cancer Center Support Grant (NCI CA060553). This research was supported in part through the computational resources and staff contributions provided by the Genomics Compute Cluster which is jointly supported by the Feinberg School of Medicine, the Center for Genetic Medicine, and Feinberg’s Department of Biochemistry and Molecular Genetics, the Office of the Provost, the Office for Research, and Northwestern Information Technology. The Genomics Compute Cluster is part of Quest, Northwestern University’s high performance computing facility, with the purpose to advance research in genomics. Graphic design was created with BioRender.com. Kishore R. Anekalla was supported by NIH T32 HL076139. Shang-Yang Chen was supported by a pre-doctoral AHA award (19PRE34380200). Alexander V. Misharin was supported by N.I.H. grants U19AI135964, P01AG049665, R01HL153312, and NUCATS COVID-19 Rapid Response Grant. Karen M. Ridge was supported by N.I.H. grants P01AG049665 and P01GM096971. Bria M. Coates was supported by funds provided by the Manne Research Institute COVID-19 Springboard Exploratory Research Award and N.I.H. grant K08HL143127. Deborah R. Winter ‘was supported by the American Lung Association, Arthritis National Research Foundation, American Thoracic Society, Scleroderma Foundation, American Federation for Aging Research, American Heart Association 18CDA34110224, and NIH/NIAID U19AI135964. G.R.S.B. was supported by NIH grants U19AI135964, P01AG049665, P01AG04966506S1, R01HL147575 and Veterans Affairs grant I01CX001777.

## Author contributions

C.M.K., K.R.A., Y.H., B.M.C., A.V.M., D.R.W., K.M.R., contributed to the acquisition, analysis, and interpretation of the data and drafting the manuscript. M.C., G.G., S.C., M.T., Y.C., J.M.D., contributed to the acquisition and interpretation of data. C.M.K., K.R.A., Y.H., B.M.C., H.A.V., P.A.R., A.V.M., G.R.S.B., D.R.W., K.M.R. contributed to the conception and design of the study and interpretation of the data. All authors critically revised and approved the final version of the manuscript.

## Figure Legends

**Supplemental Figure 1.**
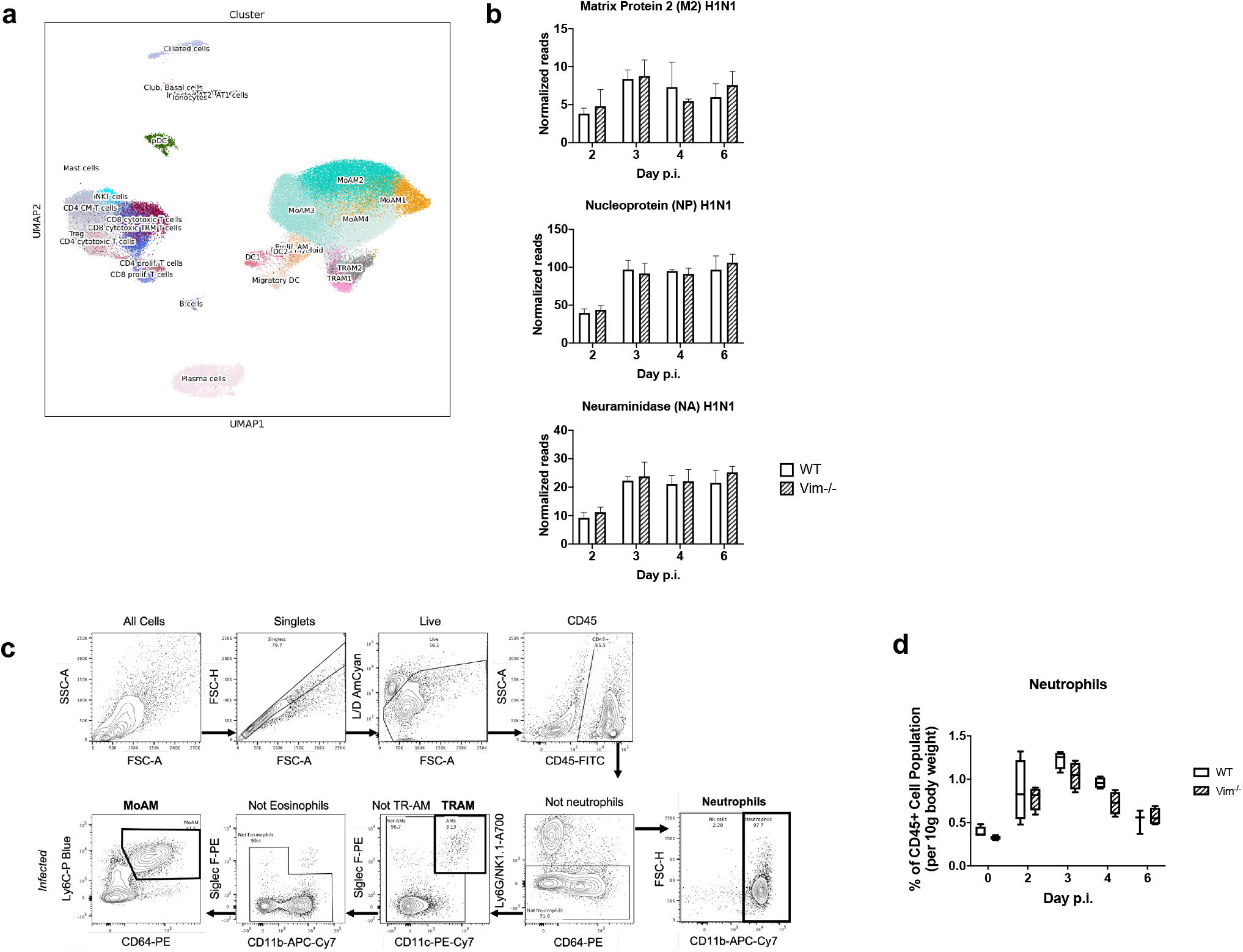
**a**) UMAP of 77,146 cells annotated by cell type obtained from 10 patients with severe COVID-19 disease within 48 hours after intubation. **b**) Quantification of H1N1 viral RNA molecules detected among RNA isolated from MoAMs, normalized to the number of reads uniquely aligned to the mouse genome (mm10) for each sample. Bar graphs represent mean, error bars represent SD. **c**) Gating strategy used for flow cytometry analysis of neutrophils, TRAMs, interstitial macrophages (IM), and MoAMs. **d**) Flow cytometric analysis demonstrated no differences in the number of neutrophils out of the total lung CD45+ cell population normalized to body weight between *Vim^-/-^* and WT mice during the IAV response.

**Supplemental Figure 2.**
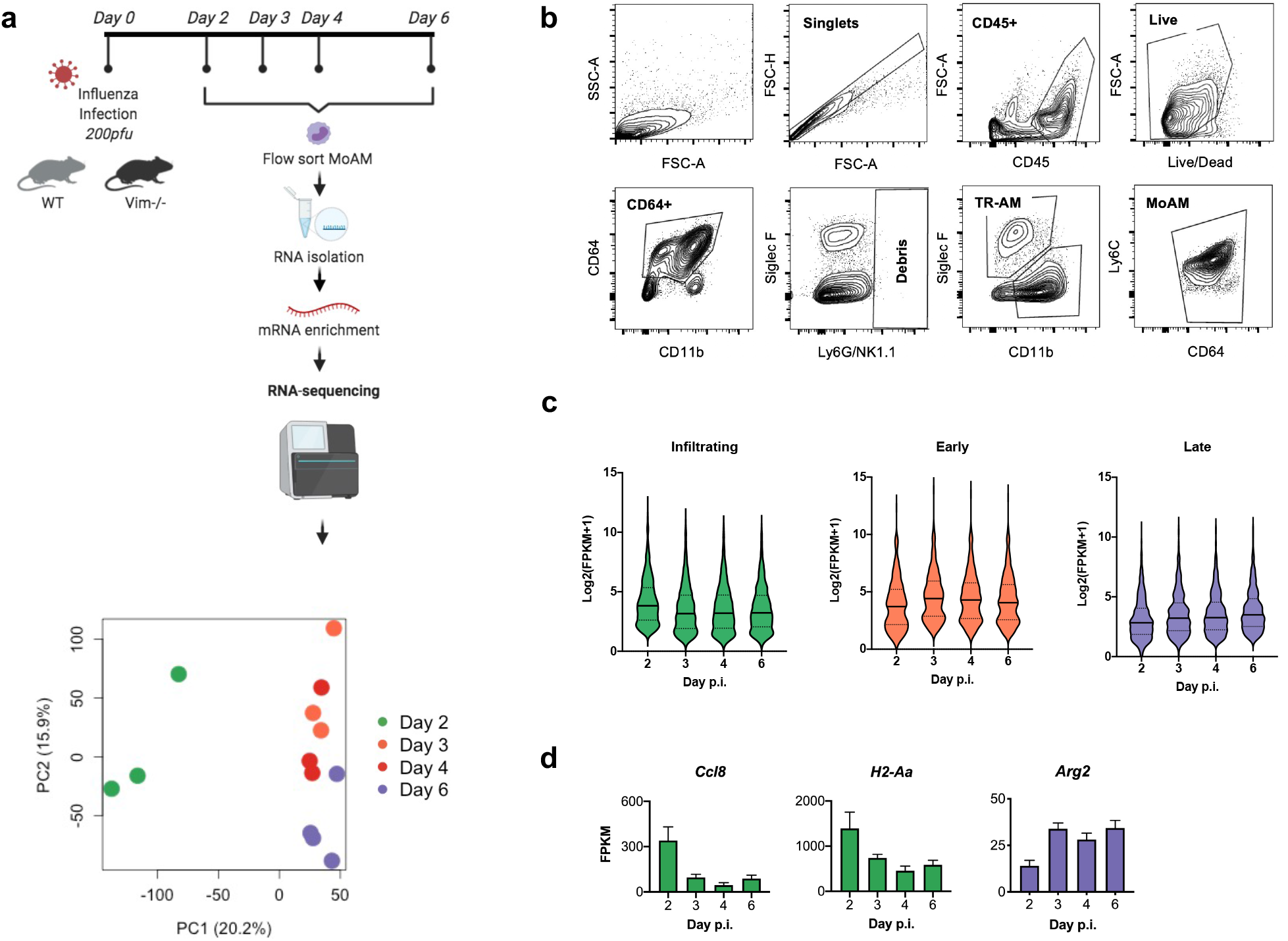
**a**) Schematic of experimental design and workflow. WT or *Vim^-/-^* mice were infected with a lethal dose of influenza A virus (200 pfu). MoAMs were sorted by flow cytometry (FACS) from lungs on 2, 3, 4, and 6 days p.i. Total RNA was isolated, mRNA was enriched, and bulk RNA-seq was performed. Principal component analysis (PCA) was used to visualize replicates. **b**) Sequential gating strategy used for FACS collection of TRAMs and MoAMs **c**) Violin plots of log_2_(FPKM+1) counts data for all genes in each cluster. Solid lines represent median, dotted lines represent quartiles. **d**) Bar graphs of gene expression (FPKM) in MoAMs for representative genes in each phase. Error bars represent SD. (ANOVA p-value <0.05).

**Supplemental Figure 3.**
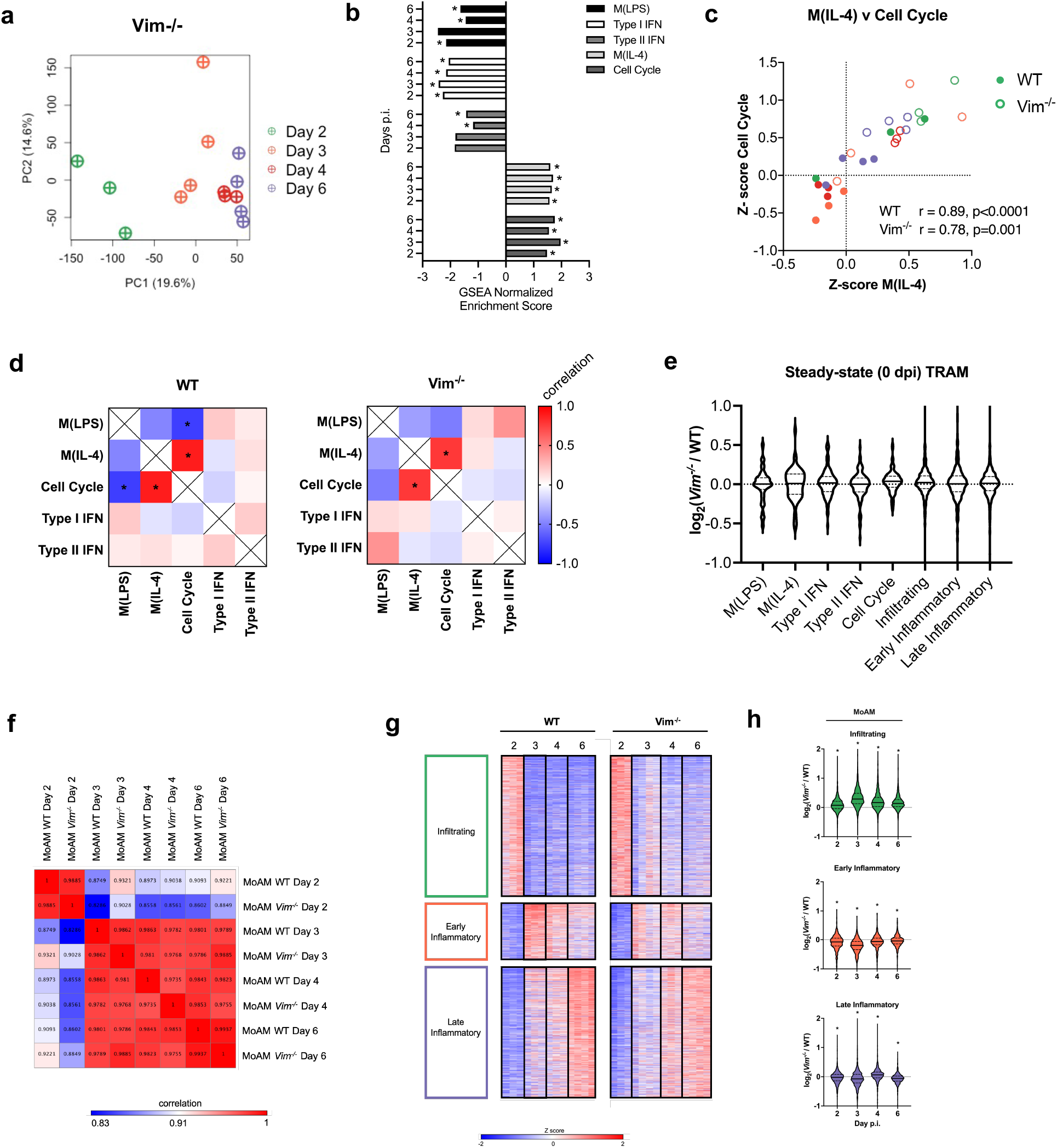
**a**) Principal component analysis (PCA) of *Vim^-/-^* MoAMs replicates. **b**) Gene Set Enrichment Analysis (GSEA) enrichment plots. GSEA was performed on gene lists ranked by edgeR reported fold-change between *Vim^-/-^* and WT MoAMs at each time point. Normalized enrichment scores (NES) are shown. Asterisks indicate statistically significant adjusted p-values <0.05. **c**) Scatter plot of z-score normalized gene expression across genes in M_(IL-4)_ vs Cell Cycle. Pearson correlation coefficients (r) across WT or *Vim^-/-^* MoAM replicates, and associated p-values are shown. **d**) Heatmap matrix of Pearson correlation between processes, asterisk indicates p-value <0.05. **e**) Violin plots representing the log_2_FC between *Vim^-/-^* and WT in tissue-resident alveolar macrophages (TRAMs) at steady-state (0 dpi) for each gene by biological cluster and biological process. Solid lines represent median, dotted lines represent quartiles **f**) Heatmap showing pairwise Pearson correlation of mean gene expression across replicates between each time point in WT and *Vim^-/-^* MoAMs. Pearson correlation coefficient values are shown within each square. Scale bar represents Pearson correlation coefficient. **g**) Heatmap of normalized expression of 2,623 genes as defined in Figure 2a. *Vim^-/-^* data was normalized to the WT mean and standard deviation (adapted z-score). **h**) Violin plots representing the log2FC between *Vim^-/-^* and WT for each gene by cluster in MoAMs. Solid lines represent median, dotted lines represent quartiles. Asterisks indicate adjusted p-value <0.05

**Supplemental Figure 4.**
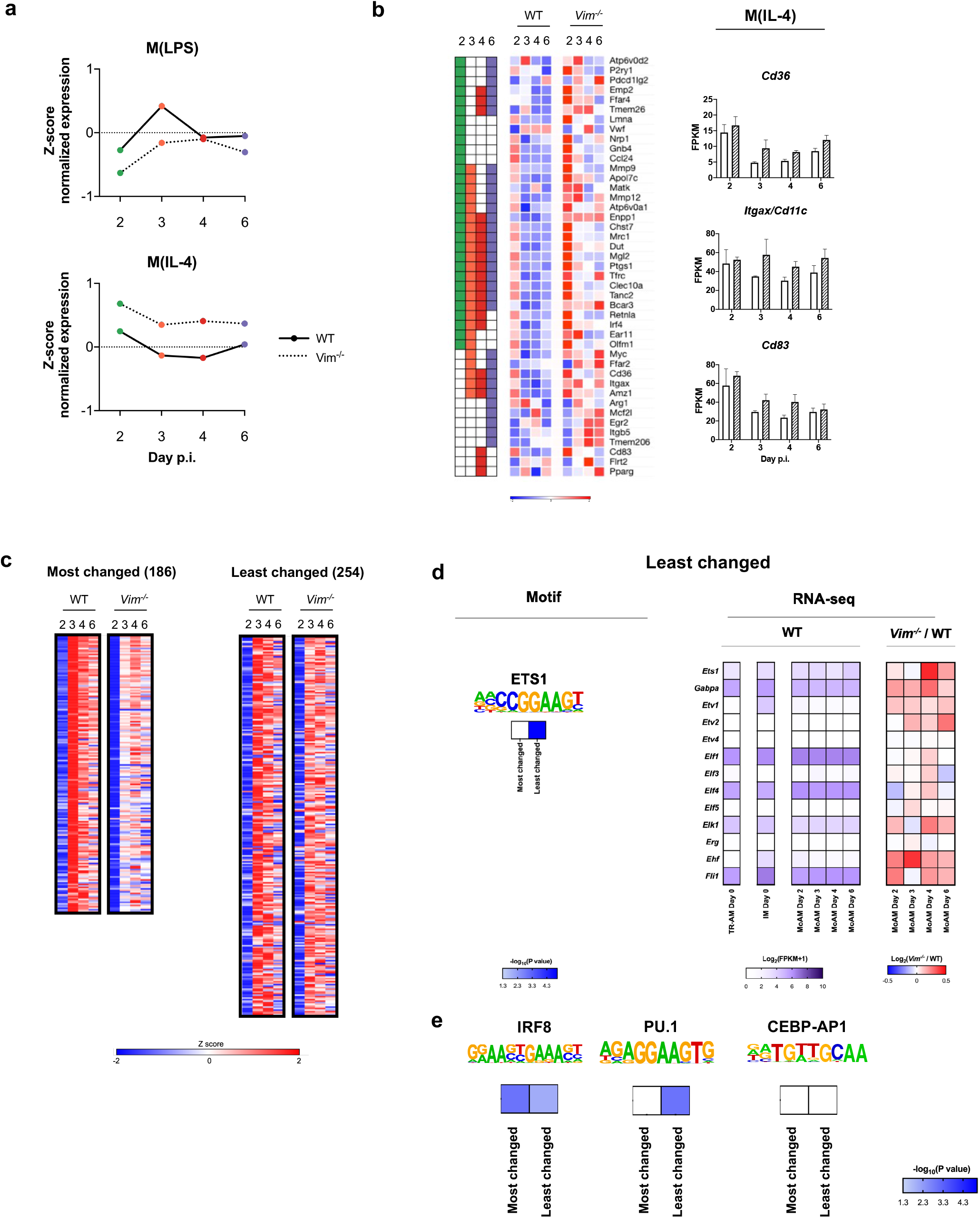
**a**) Mean adapted z-scores for M_(LPS)_ and M_(IL-4)_ genes present in the leading edge of at least 1 day from GSEA in WT and *Vim^-/-^* MoAMs. **b**) Heatmap of normalized expression of M_(IL-4)_ genes present in the leading edge of at least 1 day from GSEA. Adapted Z-score normalization for each row was applied to gene expression (FPKM) data. Bar graphs of gene expression (FPKM) for representative M_(IL-4)_ genes present in the leading edge of at least 1 day from GSEA. Error bars represent SD. **c**) Heatmap of normalized expression of 186 most changed, and 254 least changed genes on 3dpi of the Early Inflammatory Phase. Adapted Z-score normalization for each row was applied to gene expression (FPKM) data. **d**) HOMER known motif analysis using genomic background of least changed genes on 3 dpi in the Early Inflammatory Phase (254 genes). Motif logos are shown below motif names. Log_2_(FPKM+1) average gene expression of transcription factor family members is shown for WT TRAMs and WT IMs at steady-state (0 dpi) and 2-6 dpi for WT MoAMs. log2FC between *Vim^-/-^* and WT MoAM gene expression of transcription factor family members on each dpi. (n=3-4). **e**) Motif logos are shown below motif names. −log_10_pvalues of enrichment in most or least changed gene set is shown.

**Supplemental Figure 5.**
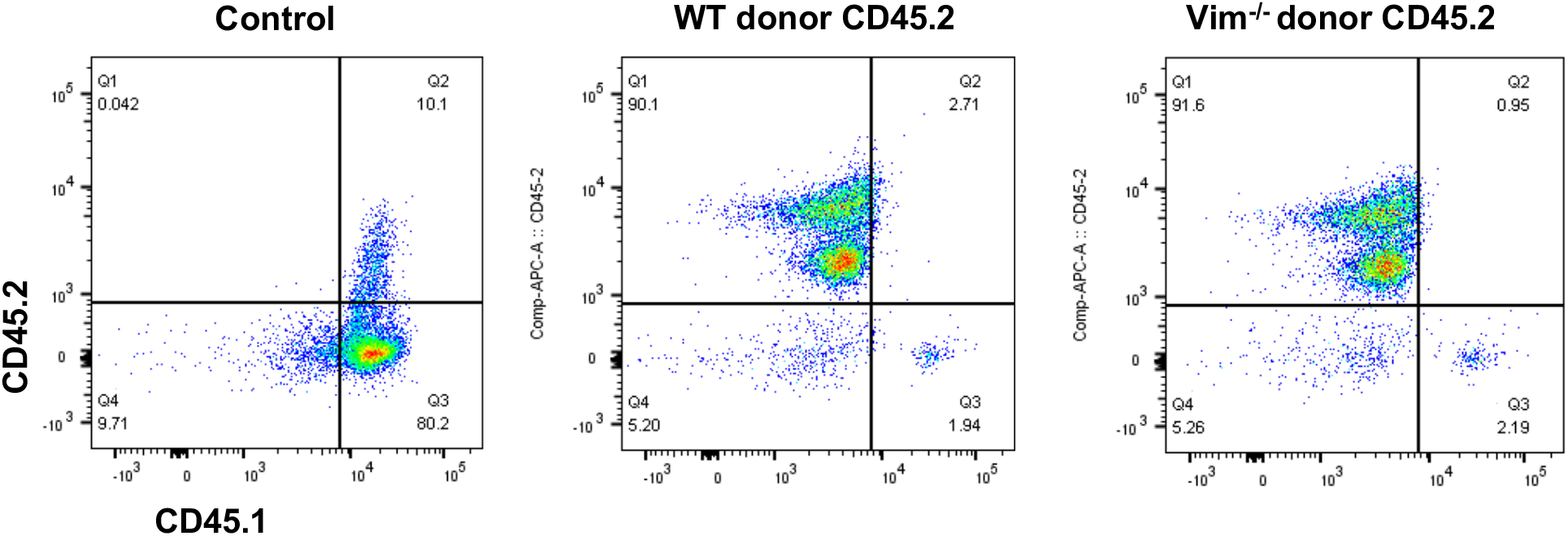
Representative flow cytometry dot plots demonstrate successful engraftment of WT and *Vim^-/-^* CD45.2 donor cells into CD45.1 recipient mice.

## Notes

### Competing Interest Statement

The authors have declared no competing interest.

